# Machine Learning Analysis to Define Cell Lineage in Leiomyosarcoma

**DOI:** 10.64898/2026.05.08.723931

**Authors:** David G.P. van IJzendoorn, Joanna Przybyl, Trevor Hastie, Judith V.M.G. Bovee, Magdalena Matusiak, Matt van de Rijn

## Abstract

**Introduction:** Cellular differentiation and lineage commitment are known to be associated with differences in DNA methylation. Leiomyosarcoma (LMS) is a tumor thought to originate from smooth muscle cells in the walls of vessels in the soft tissue (STLMS) or from the uterine myometrium (ULMS). Here, we identify the methylation signatures of normal smooth muscle cells from blood vessels and the uterine wall and compare these with those found in STLMS and ULMS. We hypothesized that these methylation signatures could be used to assign a smooth muscle subtype of origin to individual leiomyosarcomas, and that tumors of different origin would show biological differences with potential therapeutic relevance.

**Methods:** To define methylation profiles for smooth muscle from vessel walls versus those found in myometrium, EPIC methylation profiling was performed on DNA from 49 formalin-fixed paraffin-embedded (FFPE) normal smooth muscle samples. A supervised machine learning algorithm (Random Forest) was used to distinguish the methylation patterns of normal smooth muscle cells in vessel walls from those in the myometrium. The resulting classifier was applied to methylation data on 67 cases of LMS with corresponding bulk RNAseq data to identify which tumors showed a methylation signature most consistent with either blood vessel wall (LMS^vessel^) or myometrial smooth muscle (LMS^wall^). A custom signature matrix derived from scRNAseq data from 6 samples of LMS was used in CIBERSORTx analysis to compare the cellular composition of LMS cases with a vessel or uterine wall methylation signature.

**Results:** A high degree of correlation was found between the known site of origin for LMS (STLMS vs ULMS) and the methylation signature derived from different types of normal smooth muscle. LMS^wall^ tumors compared to LMS^vessel^ tumors had significantly higher activation of the PD-1 checkpoint pathway in RNAseq analysis. Digital flow cytometry by CIBERSORTx analysis showed an increased expression of transcriptomic signatures of several immune cell subtypes in LMS^vessel^ tumors.

**Conclusion:** Using a supervised machine learning approach we classified LMS samples as either showing a high similarity in methylation patterns to normal smooth muscle cells of either the vessel wall or the myometrium. We found a correlation between LMS showing either a “vessel” or “muscle wall” methylation signature and their site of origin, but notably we also identified some exceptions. When classified based on their methylation signature LMS^wall^ and LMS^vessel^ differed in their PD-1 pathway activation and in their predicted immune cell populations, suggesting potential implications for immunotherapeutic approaches.

## Introduction

LMS is a malignancy that is thought to originate from smooth muscle cells and it is hypothesized that cases of LMS found in soft tissue (STLMS) originate from the smooth muscle cells in the wall of blood vessels while those arising in the uterus (ULMS) originate from smooth muscle cells in the wall of the uterus. Here we investigate to which extent the differentiation state of normal smooth muscle cells is maintained in the LMS arising in different sites.

Cellular differentiation and lineage commitment are associated with distinct DNA methylation patterns, and paraffin-embedded specimens represent an excellent source for DNA isolation and methylation profiling from precisely defined macrodissected areas within the tissue. Furthermore, methylation profiling enables data integration with external datasets such as those from The Cancer Genome Atlas (TCGA).

In a novel approach we decided to study the lineage of smooth muscle in LMS samples using methylation profiling. We first detected significant differences in the methylation patterns of smooth muscle cells from normal vessel wall vs. myometrium, and a classifier was trained to identify the signature of smooth muscle lineage. Application of this classifier to 67 uterine and extra-uterine LMS cases matched the majority of these tumors to their site of origin. However, a subset of samples showed a methylation signature discordant with their anatomical site of origin: ULMS cases with a vessel-type signature, and STLMS cases with a myometrial-type signature.This research provides insights into the pathogenesis and heterogeneity of LMS, and suggests that recognition of, for example, a uterine LMS with a signature indicating vessel lineage may have important consequences for treatment. We further show that LMS^vessel^ and LMS^wall^ tumors differ in their immune microenvironment, with LMS^vessel^ cases displaying higher predicted immune cell infiltration and LMS^wall^ cases showing greater PD-1 pathway activation; differences that may have implications for immunotherapy stratification.

## Materials & Methods

### Normal smooth muscle and LMS samples

Normal smooth muscle samples were obtained from surgical specimens resected for unrelated clinical conditions. FFPE blocks from LMS specimens were obtained from the LMSDR Tissue Bank with institutional review board approval (Stanford University IRB-22930). For smooth muscle cells found in vessel walls 22 specimens were used from hilar liver, pulmonary and renal vessels. Myometrial smooth muscle cells were isolated from 14 uterine specimens. A further 13 samples were included from the wall of the large intestine and bladder. Archival LMS samples from 67 patients with known clinical follow-up were obtained through a collaboration with the Leiomyosarcoma Direct Research Foundation (LMSDR), a patient organization focused on LMS patients. Clinical follow-up data are available (supplementary table 1). The LMSDR cohort included 22 soft tissue LMS (STLMS) and 45 uterine LMS (ULMS) patients with follow-up data (table 1). Detailed patient characteristics are provided in supplementary table 1.

**Table 1.**
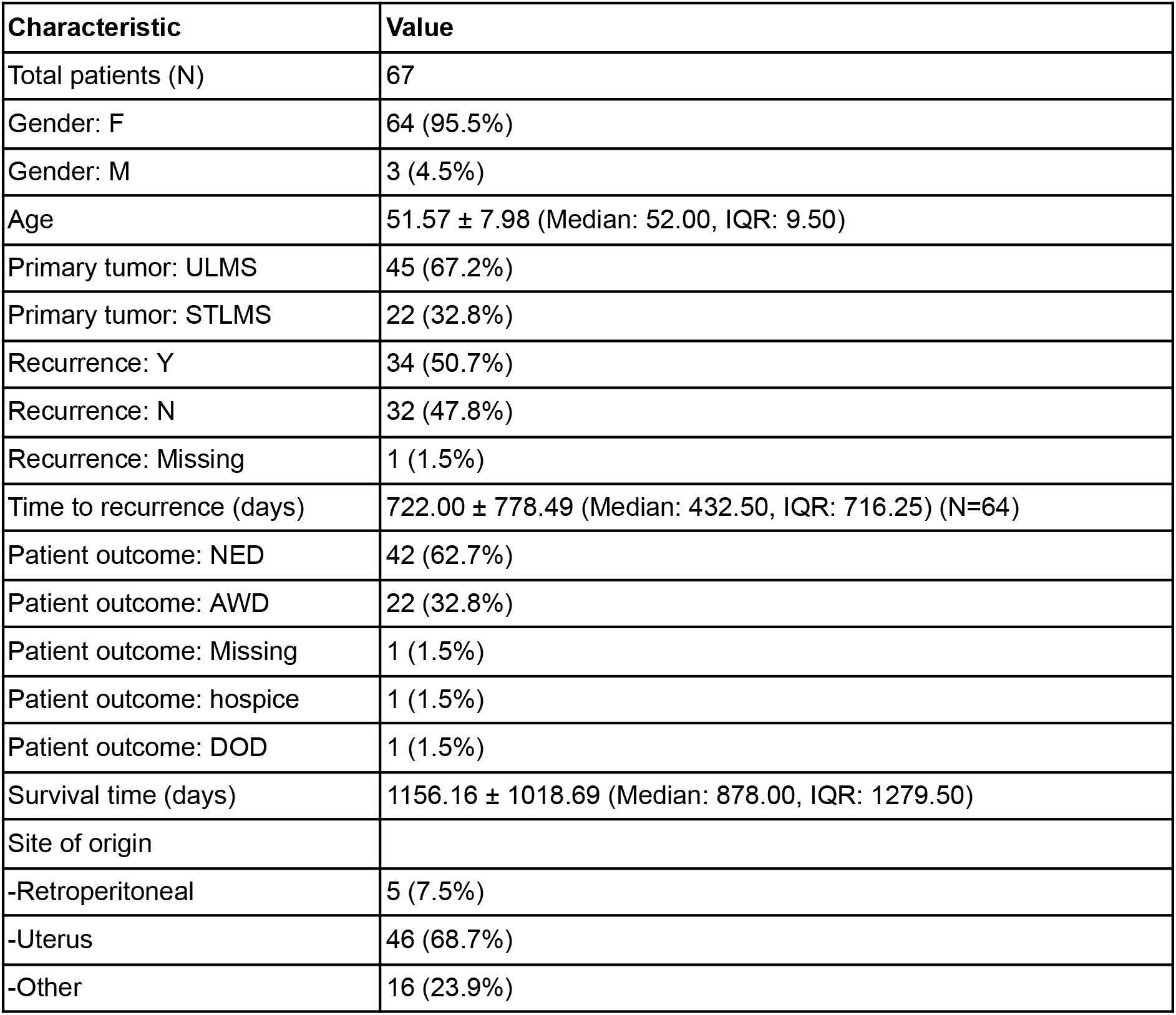
Summary of patient characteristics.

### EPIC methylation profiling and data normalisation

EPIC methylation profiling was performed on 49 samples of normal smooth muscle including 22 samples of vascular walls and 27 samples of organ smooth muscle wall (14 myometrium samples and 13 samples from smooth muscle of other hollow organs). For smooth muscle found in vessel walls, sections of vascular margins of tumor resections were used while for the wall muscle samples care was taken to select blocks containing hollow organ smooth muscle without large vessels. Multiple 2 mm-diameter core punch biopsies were taken from paraffin blocks to ensure normal smooth muscle and tumor cell purity. Smooth muscle was isolated from cases unrelated to the leiomyosarcoma samples. DNA was extracted from 2mm tissue cores using AllPrep DNA/RNA FFPE Kit, including RNase A treatment, according to the manufacturer’s protocol (Qiagen). DNA methylation profiling was performed using the Illumina Infinium MethylationEPIC BeadChip array as previously described^1^. Raw data were processed using a custom R pipeline with quality control requiring >90% of probes detected at P < 0.05. Data normalization was performed using Illumina preprocessing (preprocessIllumina function, minfi package), and poor-quality probes, SNP-associated probes, cross-reactive probes, and sex chromosome probes were removed. B and M values are available through a public GitHub repository (https://github.com/davidvi/LMSDR). Methylation samples were additionally analyzed using the Heidelberg sarcoma classifier (Koelsche et al. 2021) by uploading raw IDAT files to the Epignostix web portal (https://app.epignostix.com).

### FFPE bulk RNA sequencing

RNA was extracted in parallel with DNA extraction from the same leiomyosarcoma tissue cores using AllPrep DNA/RNA FFPE Kit, including DNase treatment, according to the manufacturer’s protocol (Qiagen). RNAseq libraries were constructed for next-generation sequencing-based expression profiling using Kapa RNA Stranded with RiboErase Library Prep Kit (Roche). Sequencing was conducted on the NovaSeq platform (Illumina) in PE150 mode. Sequencing reads were mapped to the reference genome hg38 using STAR, and the transcripts were assigned to protein coding genes using featureCounts function (Rsubread package). All expression profiling data are publicly accessible through the Gene Expression Omnibus (GEO) under accession numbers GSE315688.

### Tumor dissociation and single cell RNA profiling

We previously described our dissociation and sequencing protocol^2^. In brief, approximately 250 mg of tissue was minced and dissociated using the Tumor Dissociation Kit (130-095-929; Miltenyi Biotec) with a GentleMACS Octo Dissociator equipped with heaters (130-095-937; Miltenyi Biotec). Sample quality was assessed with an H&E stain on a smear, and cell numbers were quantified using the TC10 cell counter (BioRad). GEMs were prepared following the Chromium GEM protocol (10xGenomics). Sequencing was conducted on the NovaSeq6000 platform (Illumina) using an S1 Flow Cell (Illumina). The samples were processed using the Cell Ranger workflow (10X Genomics), and the resulting data was further analyzed using Seurat 5.0 in R. All single cell sequencing data is publicly accessible through the GEO under accession number GSE315689.

### Machine learning analysis

Two supervised DNA methylation classifiers were trained on the smooth muscle samples: a Random Forest classifier and a Support Vector Machine (SVM). Both classifiers were implemented in Python v3.11 using the Scikit-learn package. For the SVM classifier the default settings were used. For the Random Forest classifier the number of trees was set to 20 000. Data was split into a test and training set, and the top 10% most variable features (CpGs) from the full dataset were used to train the classifiers. Both classifiers reached an accuracy of 1.0 on the test data. The different smooth muscle subtypes were equally represented in the train and test datasets.

### Cluster analysis of RNA-seq data

ConsensusClusterPlus was used as previously described to perform unsupervised clustering analysis on the bulk RNA-seq samples ^3^. Similar settings were used on the current data; the analysis was run over 1,000 iterations with the following settings: distance metric of (1-Pearson correlation), 80% sample resampling, 80% gene resampling, maximum evaluated k of 12, and agglomerative hierarchical clustering algorithm. Identified clusters were compared to the previously identified clusters (as defined by Guo et al) by calculating the Jaccard index based on the differentially expressed genes for each cluster.

### Single cell RNA-seq data analysis

Seurat (v5.0) ^4^ was used in R (v4.40) to normalize and annotate the raw single-cell RNA-seq data from LMS samples. The Seurat best practices vignette for data processing was followed. Briefly, cells with low or excessively high numbers of features were removed (fewer than 200 or more than 2,500 features, respectively). Cells with high mitochondrial gene content were removed using sample-specific cut-offs. Samples were integrated, and dead cells and doublet-containing clusters were manually removed. Cell clusters were annotated based on canonical marker genes.

### CIBERSORTx digital flow cytometry analysis

CIBERSORTx was used for digital flow cytometry analysis of bulk RNA-seq samples ^5^. A custom deconvolution signature matrix was created based on the annotated single cell RNA-seq dataset containing LMS samples to define LMS-specific gene expression signatures of the different cell types in the tumor micro environment, including neoplastic cells, immune cells and stromal cells. This matrix was used in CIBERSORTx to determine cell type fractions for the 67 bulk RNA-seq LMS samples. Data was further analysed using R (v3.40). The signature matrix is publically available through GitHub (https://github.com/davidvi/LMSDR).

### Bioinformatics analysis and visualisation

For data analysis both R (v3.40) and Python (v3.11) were used. Figures were generated using either Matplotlib (v3.7.1) or ggplot2 (v3.5.2). Principal Component Analysis was performed using scikit-learn (v1.3.0). Differential expression analysis was performed using DESeq2 (v1.48.1), using a cutoff of adjusted p-value of 0.05. For Gene Set Enrichment Analysis, clusterProfiler (V3.0.4) was used. The gseKEGG function was used to identify enriched KEGG pathways. Results were ordered by EnrichmentScore and filtered for significant adjusted P values.

## Results

### Characterization of the LMSDR cohort

The LMSDR cohort consists of LMS cases with clinical follow-up available. In previous studies we and others identified 3 distinct transcriptional subtypes of LMS ^3,6–9^. To determine whether the LMSDR cohort of 67 cases was comparable to other cohorts in the literature we used ConsensusClusterPlus, and identified three distinct transcriptional subtypes within the current LMSDR RNAseq dataset. To assess the degree of similarity between these three identified subtypes and those previously reported (Guo), we computed the Jaccard index based on the top 2000 differentially expressed genes within each cluster. This analysis revealed a significant overlap between the clusters from both studies, indicating that the current cohort showed a subtype distribution similar to the LMS cases studied in previous studies and thus appeared representative of LMS in general (figure 1a). This finding underscores the robustness of the gene expression patterns in these three subtypes and highlights a strong alignment between the respective subtypes in terms of their molecular characteristics. Principal Component Analysis (PCA) was used to cluster samples (using all features) from the LMSDR dataset to assign STLMS and ULMS cases to the transcriptional subtypes. Similar to what was found in prior studies, Cluster III was predominantly composed of LMS samples of uterine origin, while Cluster I primarily consisted of soft tissue LMS samples. In contrast, Cluster II represented a heterogeneous mix of samples from both uterine and extrauterine LMS origins (figure 1b).

**Figure 1.**
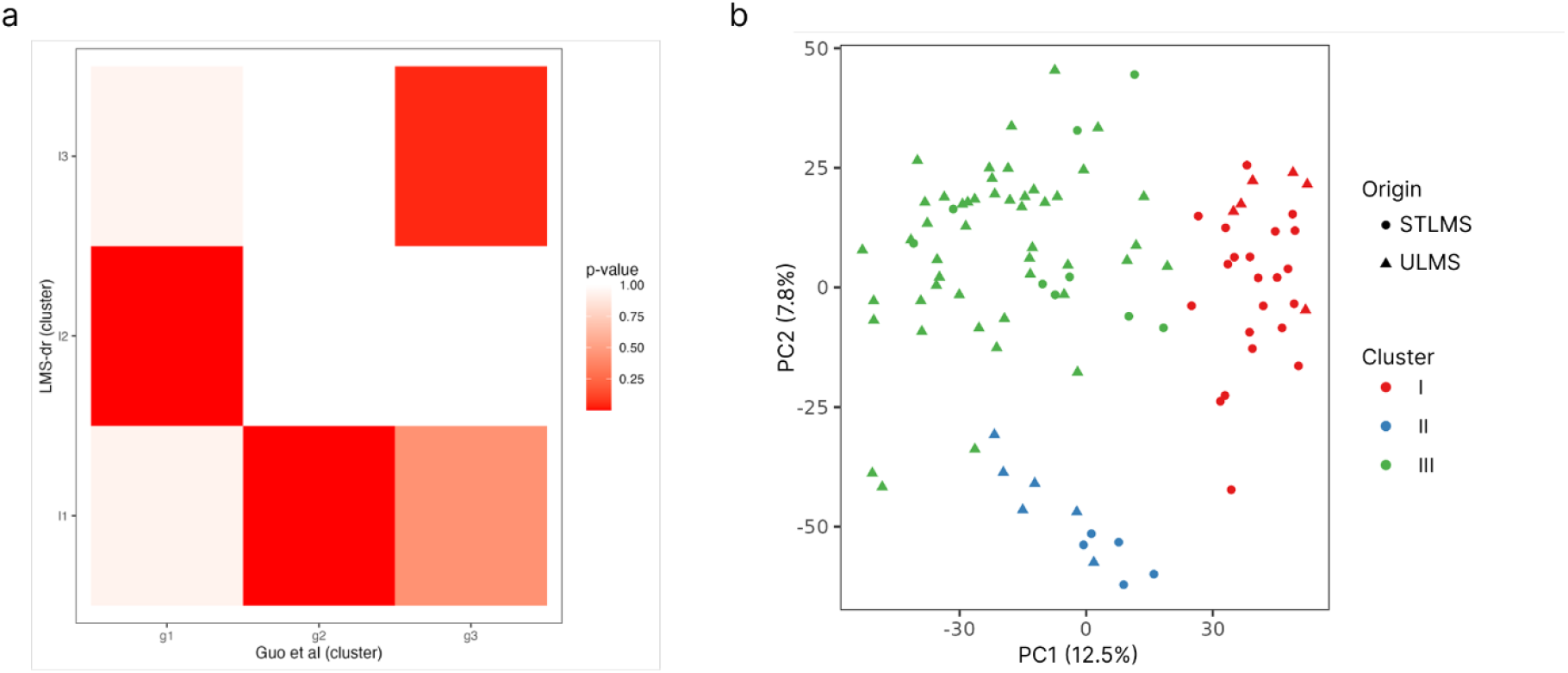
**a)** Comparison of gene expression profiles using a Jaccard index across three subtypes as described by Guo et al. and those identified in the LMSDR dataset. There exists a significant correlation among the clusters identified in both studies, indicating a strong conservation of the gene expression patterns between the respective subtypes. **b)** Principal Component Analysis (PCA) to cluster samples from the LMSDR dataset. The origin of each sample is denoted by shapes, while the respective clusters are differentiated by colors.

### Machine learning classifiers identify LMS sample similarity to vascular or myometrial smooth muscle

The EPIC arrays measured 865,859 methylation sites in each specimen, and 507,696 sites remained after quality control and removal of undetected sites. We applied a Support Vector Machine (SVM) and a RandomForest approach to determine a methylation signature that would be most distinct between cells isolated from the vessel wall or myometrium. These are two widely used machine learning algorithms that use different approaches to classify complex datasets into distinct categories. The dataset was partitioned into a training set of 32 cases and a testing set of 17 cases to validate the classifiers (figure 2a). The trained SVM and RandomForest classifiers each had an accuracy of 1.0 when run on the test dataset of 17 samples (data not shown). We next applied both classifiers to the 67 EPIC methylation profiles obtained from the LMSDR cohort and generated scores indicating the similarity of each LMS sample. All samples demonstrated highest similarity to either vascular or myometrial smooth muscle; classification as digestive system smooth muscle origin was therefore excluded from further analysis. The results from the two classifiers showed a remarkable overlap in cases classified as vessel or myometrial type methylation (now referred to as LMS^vessel^ and LMS^wall^, respectively) (figure 2b). When the methylation signature was applied to the 67 cases of LMS a pronounced overlap between the vessel-type methylation pattern and STLMS was noted, indicating a close molecular resemblance and shared differentiation state. Similarly, a substantial alignment was observed between Uterine Leiomyosarcoma (ULMS) and the myometrial smooth muscle methylation pattern.

**Figure 2.**
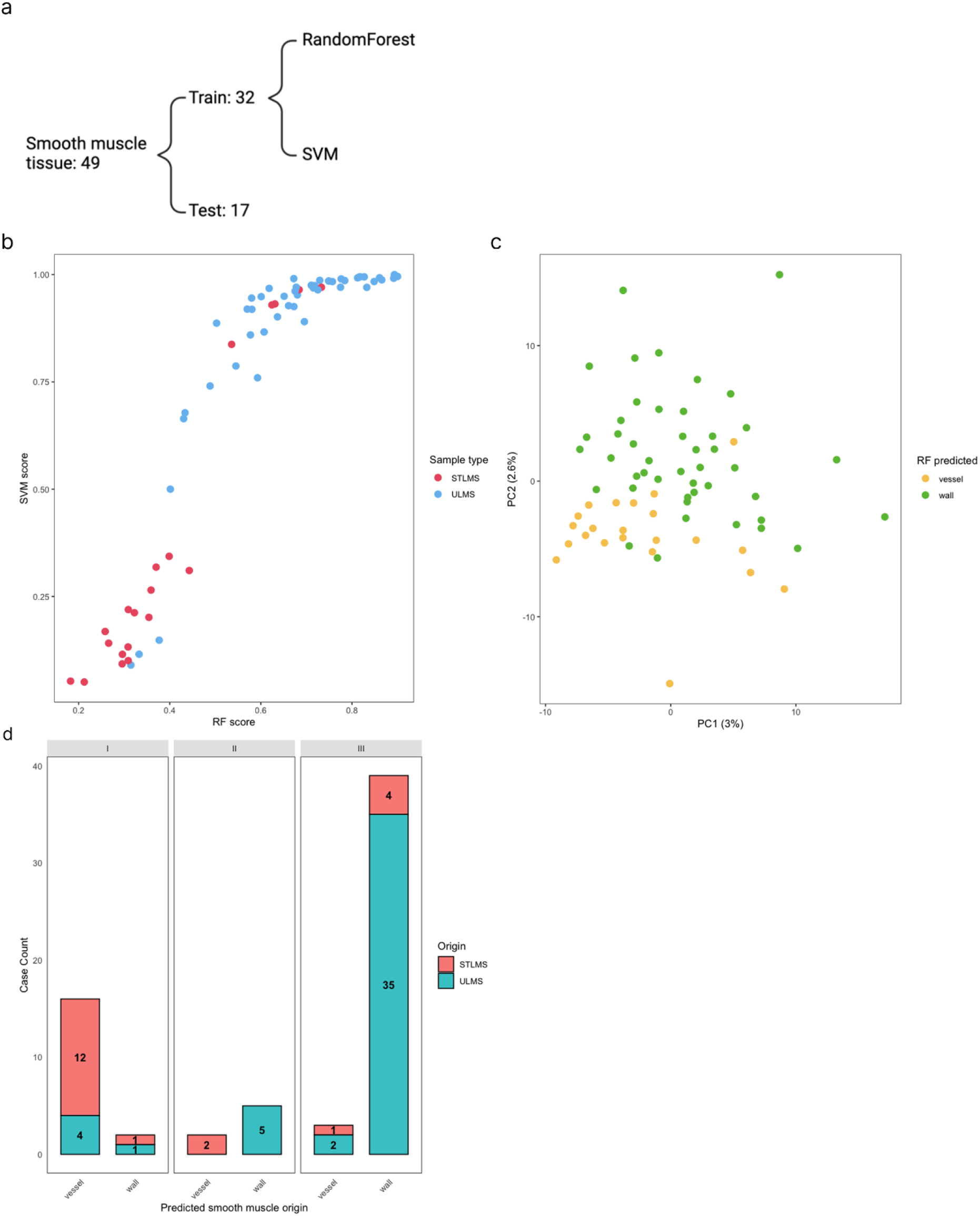
**a)** Two classifiers were generated using a Support Vector Machine (SVM) and a RandomForest. Both were trained using 49 smooth muscle samples originating from either hollow organs (13 samples), myometrium (14 samples) or vessel wall (22 samples). For the purpose of training these classifiers, the dataset comprising 49 samples was divided into training (32 samples) and testing (17 samples) sets, this split had equal representation of different types of normal smooth muscle between training and testing sets. **b)** The two classifiers were applied to the EPIC methylation profiling data on 67 LMS samples, generating scores for similarity to smooth muscle of the uterus and to smooth muscle of the vessel wall. Both classifiers showed a remarkably similar separation between the cases. Both showed a strong overlap between the methylation pattern in ULMS and normal uterine smooth muscle (high score for classifier) and between STLMS and normal vascular smooth muscle (low score for classifier). **c)** PCA analysis of the EPIC methylation profiling (using the 10% most variable features) data from the LMSDR cohort. Points are colored according to their predicted tissue of origin. LMS^vessel^ compared to LMS^wall^ shows an association with the second principal component. **d)** The LMSDR dataset was stratified by the three RNA-seq clusters, as well as by tissue of origin and Random Forest (RF) classification. This analysis revealed a strong association between methylation profiles and the previously identified RNA-seq clusters: Cluster I showed strong similarity to vessel-derived smooth muscle methylation, whereas Cluster III was closely associated with uterine smooth muscle. Cluster II appeared more evenly distributed between the two methylation profiles. The plot also highlights the existence of cases in which the RF classification does not match the anatomical site of origin.

The majority of cases demonstrated concordance between the predicted cell of origin, as indicated by the classifier, and the anatomically known site of tumor origin. However, a subset of cases lacked this correlation, suggesting that a proportion of ULMS tumors may arise from vascular smooth muscle within the uterus (7 samples, 14.9). While STLMS tumors predominantly appear to originate from vascular smooth muscle, some showed a myometrial smooth muscle signature (5 samples, 25.0%). The separation into LMS^vessel^ and LMS^wall^, based on the cell of presumed origin, revealed a compelling correlation with previously identified RNAseq clusters (figure 2c and 2d). Notably, Cluster I exhibited a strong correlation with vascular smooth muscle type of methylation (88.9%), whereas Cluster III was closely aligned with uterine smooth muscle (92.9%). This significant correlation (Fisher’s exact test, p = 6.79 × 10^−10^) underscores the relevance of these clusters in distinguishing between different LMS subtypes based on their tissue of origin.

The discordant samples were further analyzed using the Heidelberg sarcoma classifier ^10^, which includes categories for both uterine and extra-uterine leiomyosarcoma. Using the Heidelberg classifier, all 7 discordant LMS^vessel^ cases (that clinically appeared to arise in the uterus) were classified as extra-uterine leiomyosarcoma, supporting the idea that they had arisen from vessels in the myometrium. Additionally the 5 discordant LMS^wall^ cases (that clinically were thought to be extra-uterine) (table 2 and figure S1), were classified as uterine leiomyosarcoma by the Heidelberg classifier, supporting the methylation-based cell-of-origin assignments over the anatomical site of presentation. Notably, the discordant STLMS samples predominantly originated from gynaecological or perigynaecological sites (vulva, vagina, perineum, and pelvis; table 2).

**Table 2.** Overview of discordant samples.

| STT | Primary tumor | Category | RandomForest | Heidelberg |
| --- | --- | --- | --- | --- |
| 13334 | uterus | ULMS | vessel | STLMS |
| 13336 | uterus | ULMS | vessel | STLMS |
| 13357 | uterus | ULMS | vessel | STLMS |
| 13387 | uterus | ULMS | vessel | STLMS |
| 13431 | uterus | ULMS | vessel | STLMS |
| 13452 | uterus | ULMS | vessel | STLMS |
| 13464 | vulva | STLMS | wall | ULMS |
| 13477 | pelvic | STLMS | wall | ULMS |
| 13483 | uterus | ULMS | vessel | STLMS |
| 13542 | perineum | STLMS | wall | ULMS |
| 13543 | vagina | STLMS | wall | ULMS |
| 13569 | small intestine | STLMS | wall | STLMS |

Kaplan-Meier analysis between LMS^vessel^ and LMS^wall^ showed that the tissue of origin assigned based on DNA methylation classifier did not result in significant differences in recurrence-free survival within our dataset (log rank test p = 0.13). However, this might be due to the limited sample size, which may have insufficient statistical power to detect survival differences between these groups (figure S2).

### Differential analysis between the samples originating from wall and vessel smooth muscle

All samples were classified into either LMS^vessel^ or LMS^wall^ methylation type, and differential expression analysis was performed on their bulk RNA-seq transcriptional profiles (figure 3a). The LMS^vessel^ group was characterized by elevated expression of the genes encoding matrix metalloproteinases, including *MMP12* and *MMP13*, as well as immune-related genes such as *CXCL9, CX3CL1*, and *FCN1*. As expected, the LMS^wall^ group exhibited higher expression of estrogen receptor (ER)-associated genes including *ESR1* (log_2_ fold change: 3.46, p < 0.001). Progesterone receptor *PGR* (log_2_ fold change: 2.6, p < 0.001) was also upregulated in this group. KEGG pathway analysis revealed distinct pathway enrichment patterns for both groups. Notably, many pathways enriched in the LMS^vessel^ group were related to immune system activity. Among these, the PD-L1 expression and PD-1 checkpoint pathway in cancer was identified as activated in LMS^vessel^ tumors (figure 3b and figure S3). Furthermore, PD-L1 gene expression was significantly higher in the LMS^vessel^ group, and this group separation explained more of the variance in PD-L1 expression than did anatomical site of origin (STLMS vs. ULMS; adjusted R^2^ = 21.7% [random forest] vs. 8.2% [anatomical location]; figure 3c), whereas PD-L1R showed similar expression in both groups (figure S4).

**Figure 3.**
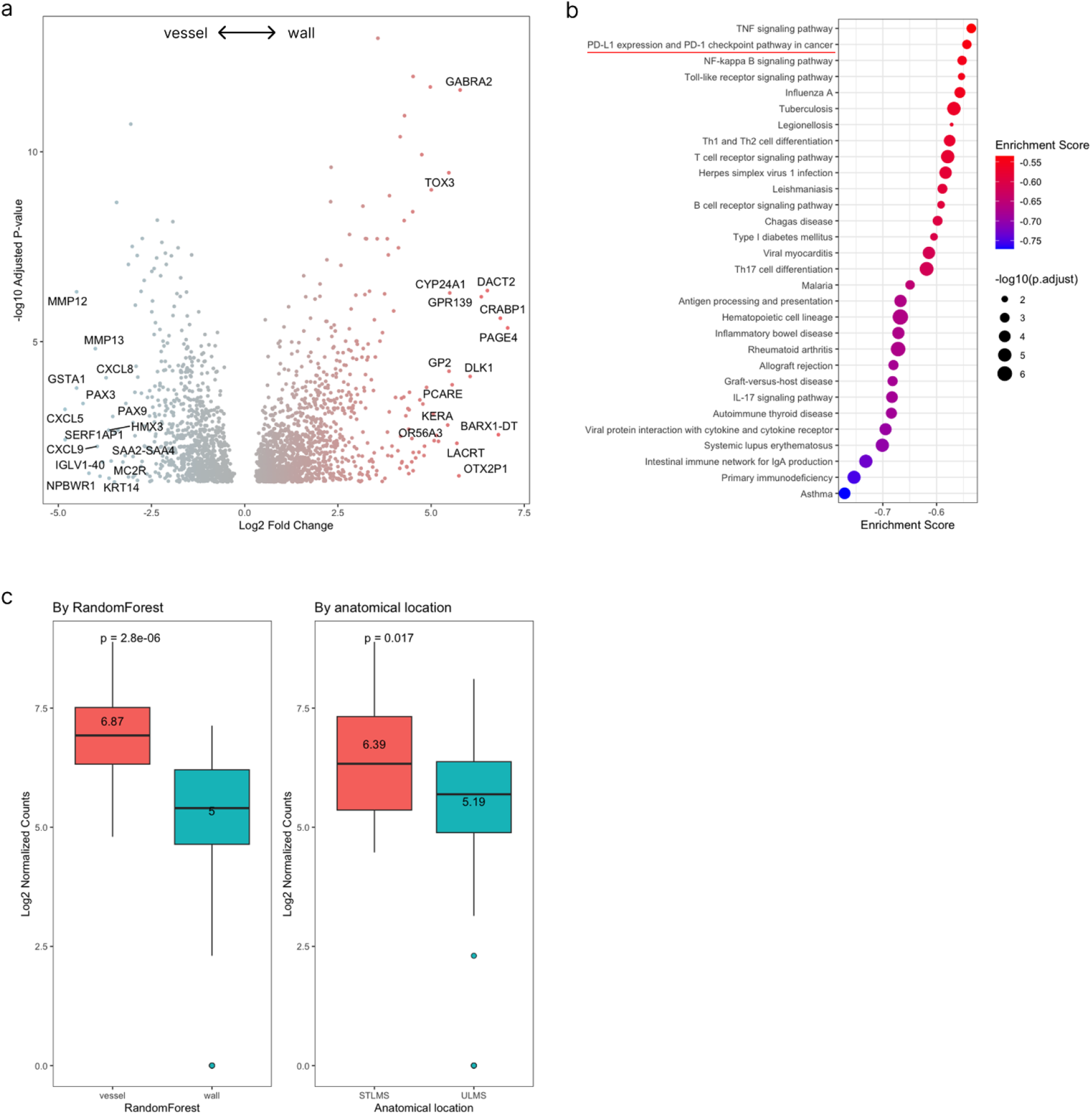
**a)** Differential expression analysis was performed on LMSDR cases, comparing samples classified as resembling smooth muscle of the uterine wall versus those resembling vascular smooth muscle. **b)** KEGG pathway analysis revealed that most significantly enriched pathways had negative enrichment scores, indicating their relative specificity to LMS samples resembling vascular smooth muscle. These included several immune-related pathways, notably the PD-1 checkpoint pathway in cancer (p.adjusted = 0.00142 and 19/22 genes are enriched). **c)** CD274 (PD-L1) expression stratified by random forest classification (LMSvessel vs. LMSwall, left) and by anatomical site of origin (STLMS vs. ULMS, right). The random forest classification separated CD274 expression more clearly than anatomical location.

### Single cell RNA-seq and CIBERSORTx identifies cellular composition of the LMS samples

We performed single cell sequencing on six LMS samples from 4 patients (not included in the methylation dataset) to generate a custom deconvolution matrix to recognize the different cell types within LMS (figure 4a). This matrix was used in CIBERSORTx to determine the percentage of different cell types in the bulk RNA-seq data for each of the 67 patient samples (digital flow cytometry) and compared the cellular composition between LMS^vessel^ and LMS^wall^ samples (detection p values are all < 0.05) (figure 4b). The findings are largely in agreement with the KEGG pathway analysis and show a statistically significant increase in expression of transcriptomic signatures of immune cell types including monocytes, CD4+ and CD8+ T-lymphocytes with a trend for increased NK cells in LMS^vessel^ cases. The deconvoluted percentage of macrophages was similar between LMS^vessel^ and LMS^wall^ and the samples with LMS^wall^-type methylation showed higher numbers of neoplastic cells (figure 4c). The analysis of the TCGA LMS sample dataset showed similar results (figure S5).

**Figure 4.**
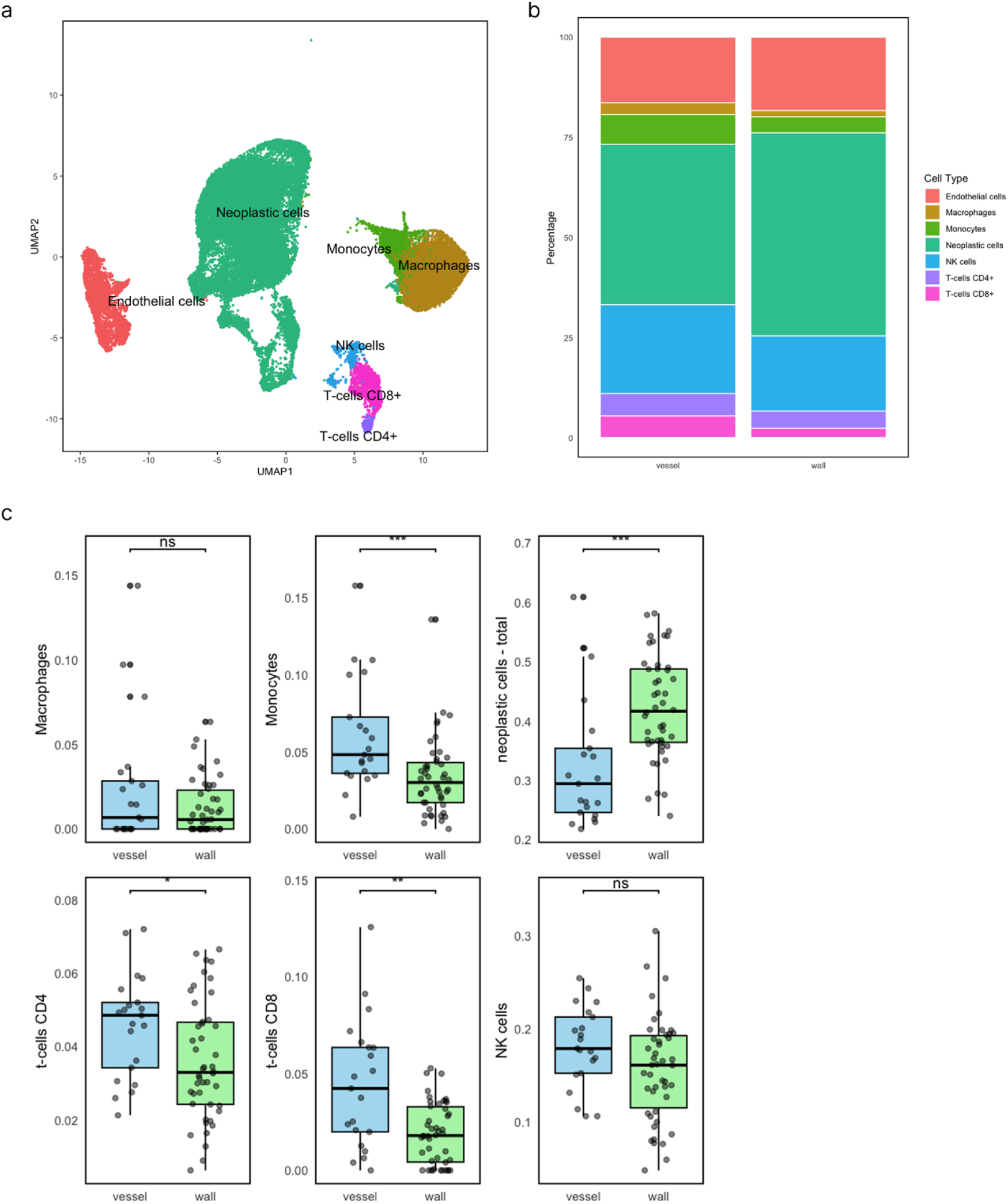
**a)** Single-cell RNA sequencing (scRNA-seq) of five LMS samples revealed distinct intertumoral clusters comprising both immune cell populations and neoplastic cell populations. **b)** Cell composition was assessed using the CIBERSORTx deconvolution matrix, comparing samples resembling vessel and wall smooth muscle. **c)** Consistent with the differential expression results, we observed an increased abundance of various immune cell types in LMS^vessel^. Significantly enriched cell types included monocytes, CD4+ T cells, and CD8+ T cells (Wilcoxon test, * = 0.05, ** = 0.01, *** = 0.001). In contrast, LMS^wall^ contained significantly more neoplastic cells.

## Discussion

In this study we investigated the smooth muscle tissue of origin for LMS. Previous studies have already identified smooth muscle as the tissue of origin for LMS, but this, to our best knowledge, has not been shown using a multi-omics approach. Our dataset included EPIC methylation profiles, bulk RNA-seq data, and single-cell RNA-seq data. In a novel approach, we performed methylation profiling on normal smooth muscle samples from vessel walls, myometrium, bladder, and colon wall, allowing us to generate a classifier that distinguishes LMS samples with a vessel wall smooth muscle signature from those with a myometrial signature.

As expected, most uterine LMS samples carried a myometrial methylation signature while soft tissue LMS samples carried a vessel wall signature. However, we also identified a subset of uterine LMS cases with a vessel wall signature and a smaller number of extra-uterine LMS cases with a myometrial signature. It is possible that the uterine cases with a signature suggesting vessel walls represent samples with a high percentage of vessels within the tumor although this was not histologically obvious in the samples used for analysis. The STLMS cases with a uterine wall signature predominantly arose from peri-uterine sites, which may account for their myometrial smooth muscle signature.

RNA-seq analysis revealed biological differences between these groups. LMS with a vessel wall signature showed enrichment for immune-related pathways, including the *PD-L1* expression and PD-1 checkpoint pathway in cancer. Digital flow cytometry based on CIBERSORTx, using an LMS-specific single-cell reference matrix, confirmed that vessel wall–type LMS harbored higher proportions of inferred immune cell types such as monocytes, CD4+ T cells, and CD8+ T cells. In contrast, LMS with a myometrial signature contained significantly more neoplastic cells. These differences were consistent across our cohort and in the TCGA LMS dataset. Consistent with our findings, prior immunohistochemical studies have reported PD-L1 expression and lymphocyte infiltration in a substantial proportion of soft tissue LMS, including the work of Kostine et al., who found PD-L1 positivity in 30% and high T-cell infiltration in 52% of a predominantly extra-uterine LMS cohort^11^.

The immune-rich phenotype of vessel wall–type LMS may have clinical relevance. Trials of immune checkpoint inhibitors in LMS have so far shown limited success, but our results raise the possibility that responses could be enriched in specific biological subsets, such as LMS of vessel wall origin. This warrants re-analysis of existing trial data to determine whether cell-of-origin signatures are associated with treatment response, and prospective evaluation in future studies. In this study, we show that *CD274* (PD-L1) expression is more strongly stratified by the classifier-predicted cell of origin than by anatomical site of presentation, suggesting that lineage-based classification may better identify candidates for checkpoint blockade. Future studies are needed to determine whether treatment decisions guided by the classifier-predicted cell lineage may be more effective than those based solely on the anatomical site of tumor origin.

We and others have previously shown that on a transcriptomic level, at least three different molecular subtypes exist within LMS^3,6–9^. Among these studies, Anderson et al. investigated the smooth muscle tissue of origin using a clustering-based analysis of transcriptome sequencing data. This approach identified three putative sites of origin: vascular, digestive, and gynecological smooth muscle. In contrast, our methylation-based approach did not identify any tumor samples with methylation profiles resembling those of digestive smooth muscle, despite the inclusion of such tissues in our reference dataset. This discrepancy may reflect fundamental methodological differences, as our classification relied on DNA methylation patterns rather than transcriptomic data.

Finally, our results reinforce prior observations of three transcriptional subtypes in LMS, as reported by Guo et al, and Anderson et al, and confirmed in our current cohort. The methylation-based classification adds an additional layer of resolution to these transcriptional subtypes, refining our understanding of LMS heterogeneity. In our cohort, cell-of-origin signatures did not correlate with clinical outcome, but larger studies will be needed to fully assess their prognostic impact.

In summary, we demonstrate that LMS can be classified according to smooth muscle cell lineage of origin, with clear molecular and immune microenvironment differences between vessel wall–derived and myometrial-derived tumors. These findings have potential implications for understanding LMS pathogenesis, refining molecular taxonomy, and guiding future therapeutic strategies.

