## Supplementary figures and images for "Machine Learning Analysis to Define Cell Lineage in Leiomyosarcoma"

### s1

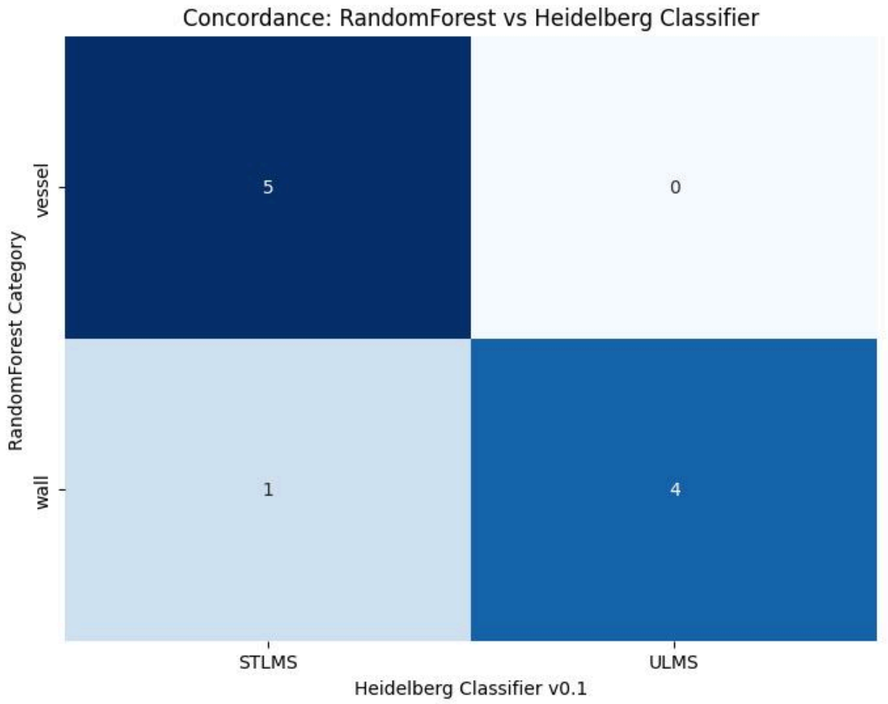

### s2

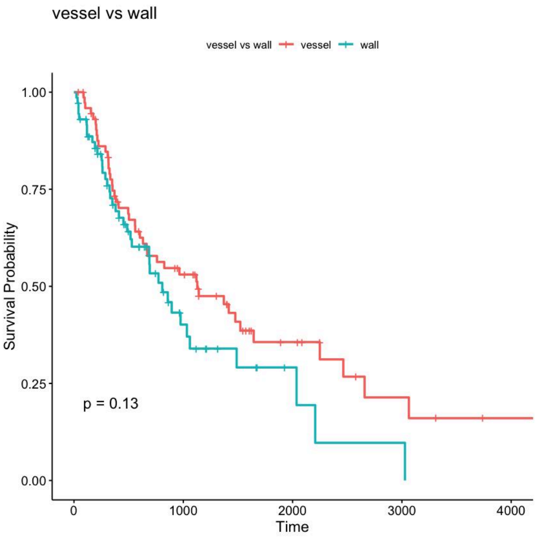

### s3

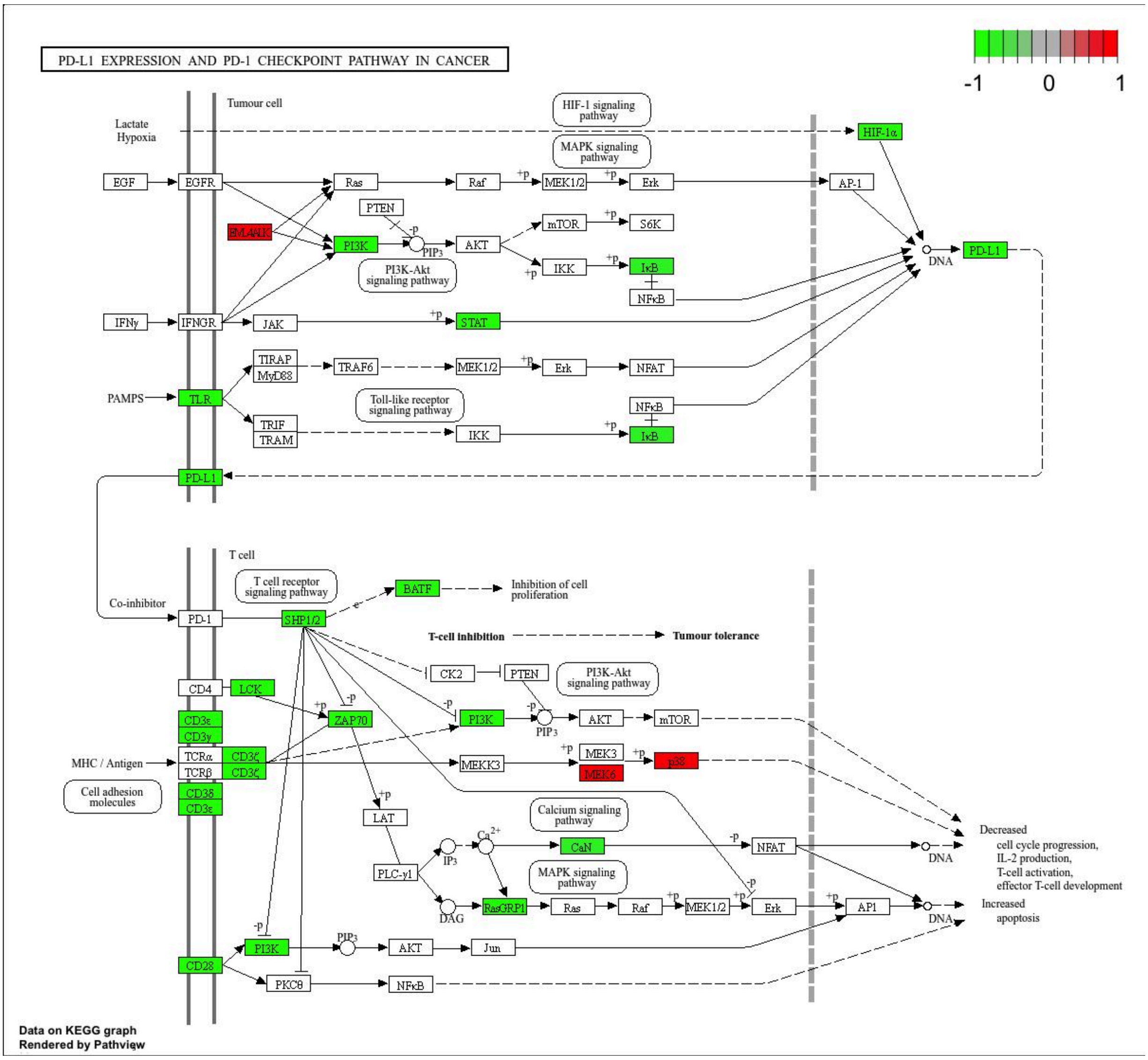

### s4

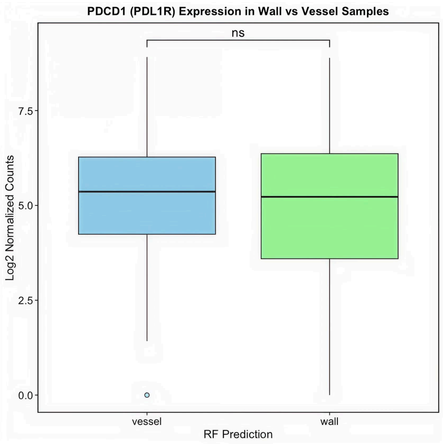

### s5

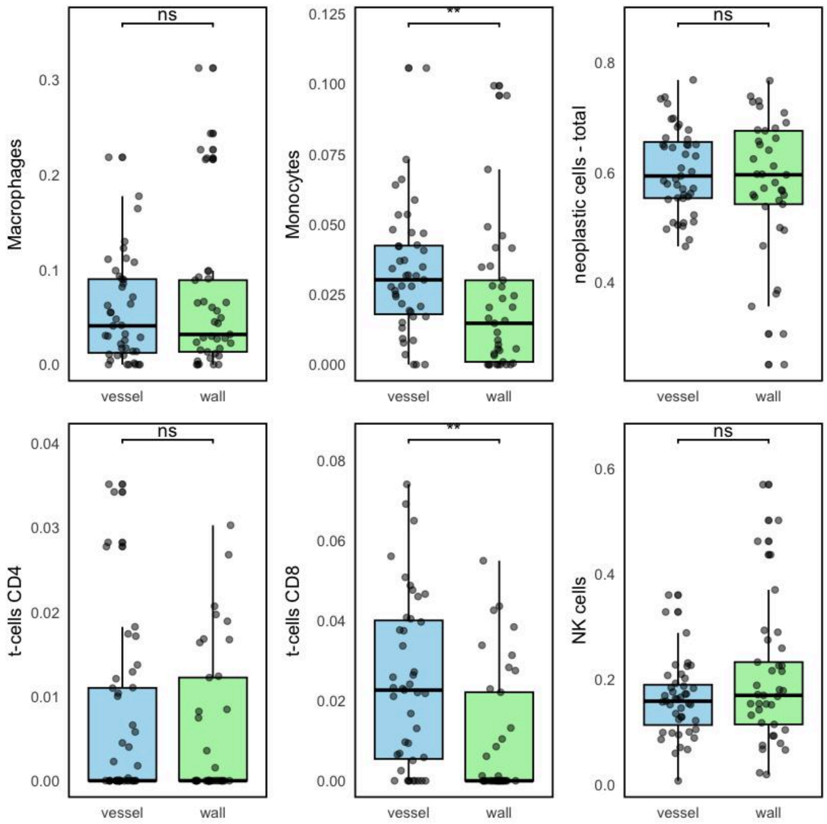
